# A conserved biosynthetic gene cluster is regulated by quorum sensing in a shipworm symbiont

**DOI:** 10.1101/2022.02.10.479910

**Authors:** Jose Miguel D. Robes, Marvin A. Altamia, Ethan G. Murdock, Gisela P. Concepcion, Margo G. Haygood, Aaron W. Puri

## Abstract

Bacterial symbionts often provide critical functions for their hosts. For example, wood-boring bivalves called shipworms rely on cellulolytic endosymbionts for wood digestion. However, how the relationship between shipworms and their bacterial symbionts is formed and maintained remains unknown. Quorum sensing (QS) often plays an important role in regulating symbiotic relationships. We identified and characterized a QS system found in *Teredinibacter* sp. strain 2052S, a gill isolate of the wood-boring shipworm *Bactronophorus* cf. *thoracites*. We determined that 2052S produces the signal *N-*decanoyl*-L*-homoserine lactone (C_10_-HSL), and that this signal controls activation of a biosynthetic gene cluster co-located in the symbiont genome that is conserved among all symbiotic *Teredinibacter* isolates. We subsequently identified extracellular metabolites associated with the QS regulon, including ones linked to the conserved biosynthetic gene cluster, using mass spectrometry-based molecular networking. Our results demonstrate that QS plays an important role in regulating secondary metabolism in this shipworm symbiont. This information provides a step towards deciphering the molecular details of the relationship between these symbionts and their hosts. Furthermore, because shipworm symbionts harbor vast yet underexplored biosynthetic potential, understanding how their secondary metabolism is regulated may aid future drug discovery efforts using these organisms.

**IMPORTANCE:** Bacteria play important roles as symbionts in animals ranging from invertebrates to humans. Despite this recognized importance, much is still unknown about the molecular details of how these relationships are formed and maintained. One of the proposed roles of shipworm symbionts is the production of bioactive secondary metabolites due to the immense biosynthetic potential found in shipworm symbiont genomes. Here, we report that a shipworm symbiont uses quorum sensing to coordinate activation of its extracellular secondary metabolism, including the transcriptional activation of a biosynthetic gene cluster that is conserved among many shipworm symbionts. This work is a first step towards linking quorum sensing, secondary metabolism, and symbiosis in wood-boring shipworms.

---

Wood-boring shipworms are bivalves that harbor intracellular gammaproteobacteria in their gills that express cellulases for wood digestion (1, 2). However, many details of the molecular mechanisms that govern the selection and maintenance of symbiotic bacteria by shipworms remain unknown. Analysis of published endosymbiont genomes and shipworm-associated metagenomes has indicated that these gill endosymbionts are also capable of producing a plethora of secondary metabolites comparable to well-known producers such as *Streptomyces* spp. (3). Several predicted biosynthetic gene clusters (BGCs) are conserved among shipworm symbionts (3), which could indicate that the products of these clusters play a role in the symbiotic relationship. For example, the boronated antibiotic tartrolon, isolated from the shipworm symbiont *Teredinibacter turnerae* T7901, is hypothesized to participate in the inhibition of competing parasites in the shipworm gills and/or cecum (4).

Bacterial symbionts often use quorum sensing (QS) to coordinate group behavior, which is thought to help differentiate between a low-density, free-living state, and high-density, host-associated state (5). In many proteobacteria, QS is mediated by acyl-homoserine lactone (acyl-HSL) signals produced by LuxI-family synthases (6). In this type of QS system, genes are regulated by members of the LuxR-family of transcription factors which bind and respond to acyl-HSLs (6). The first QS system was characterized in the invertebrate symbiont *Aliivibrio fischeri*, which uses 3-oxo-hexanoyl-*L*-homoserine lactone (3-oxo-C_6_-HSL) to regulate bioluminescence in the light organ of its host squid, *Euprymna scolopes* (7, 8). Characterization of QS systems in shipworm symbionts therefore has the potential to provide insight into the details of their relationship with their host.

QS often regulates the production of extracellular factors, including secondary metabolites and enzymes such as proteases (6, 9–11). A common example is the plant-associated pathogen, *Erwinia carotovorum*, which is known to produce the antibiotic carbapenem in response to QS (9). In many cases QS systems regulate adjacent genes in bacterial genomes, and a recent genome mining effort discovered that BGCs neighboring *luxR* homologs are widespread in proteobacteria (12). Interestingly, only a small percentage of QS-linked BGCs identified in this study were found in free-living and invertebrate-associated bacteria, while plant- and human-associated bacteria made up the majority (12).

One BGC of interest that is found in all cellulolytic shipworm symbionts isolated to date is a predicted hybrid *trans*-AT PKS-NRPS (*trans*-acyltransferase polyketide synthase-nonribosomal peptide synthetase) gene cluster termed GCF_3 (3). The product of GCF_3 has not been isolated or characterized. *Teredinibacter* sp. strain PMS-2052S.S.stab0a.01 (referred to here as 2052S) is a cellulolytic bacterial strain isolated from the gills of a specimen of the shipworm *Bactronophorus* cf. *thoracites* collected in Butuan, Agusan del Norte, Philippines. In the genome of 2052S, the GCF_3 BGC is adjacent to a predicted QS system. Determining how this BGC is regulated in a symbiont may enable the identification and characterization of its product.

In this work, we characterized the QS system used by the shipworm endosymbiont 2052S. We identified the acyl-HSL signal and linked it with its cognate synthase and receptor. We then determined that this QS system regulates the neighboring GCF_3 BGC, and used untargeted metabolomics and molecular networking to identify metabolites associated with the QS regulon including potential products of the GCF_3 BGC. To our knowledge, this is the first characterization of a shipworm endosymbiont QS system, which extends our understanding of the molecular details of this symbiosis.

## RESULTS AND DISCUSSION

### A conserved biosynthetic gene cluster in cellulolytic shipworm symbionts is adjacent to quorum sensing genes in strain 2052S

The cellulolytic strain 2052S was isolated from the gills of a specimen of the wood-boring shipworm *Bactronophorus* cf. *thoracites* (see Table S1 for strain isolation information) (3). It is likely an intracellular symbiont like other *Teridinibacter* species (1), however more studies will be needed to determine this definitively. In the genome of 2052S, the conserved BGC GCF_3 is adjacent to a *luxR*-family transcription factor gene (K256DRAFT_2894, *tbaR*) and an acyl-HSL synthase gene (K256DRAFT_2894, *tbaI*) **(Figure 1A)**. We therefore hypothesized that GCF_3 may be regulated by QS in this strain, as is true with other QS-linked BGCs in proteobacteria (10, 11). In other shipworm symbionts, GCF_3 is not adjacent to QS genes **(Figure S1)**, indicating that QS may have been lost in those isolates or gained in 2052S.

**Figure 1.**
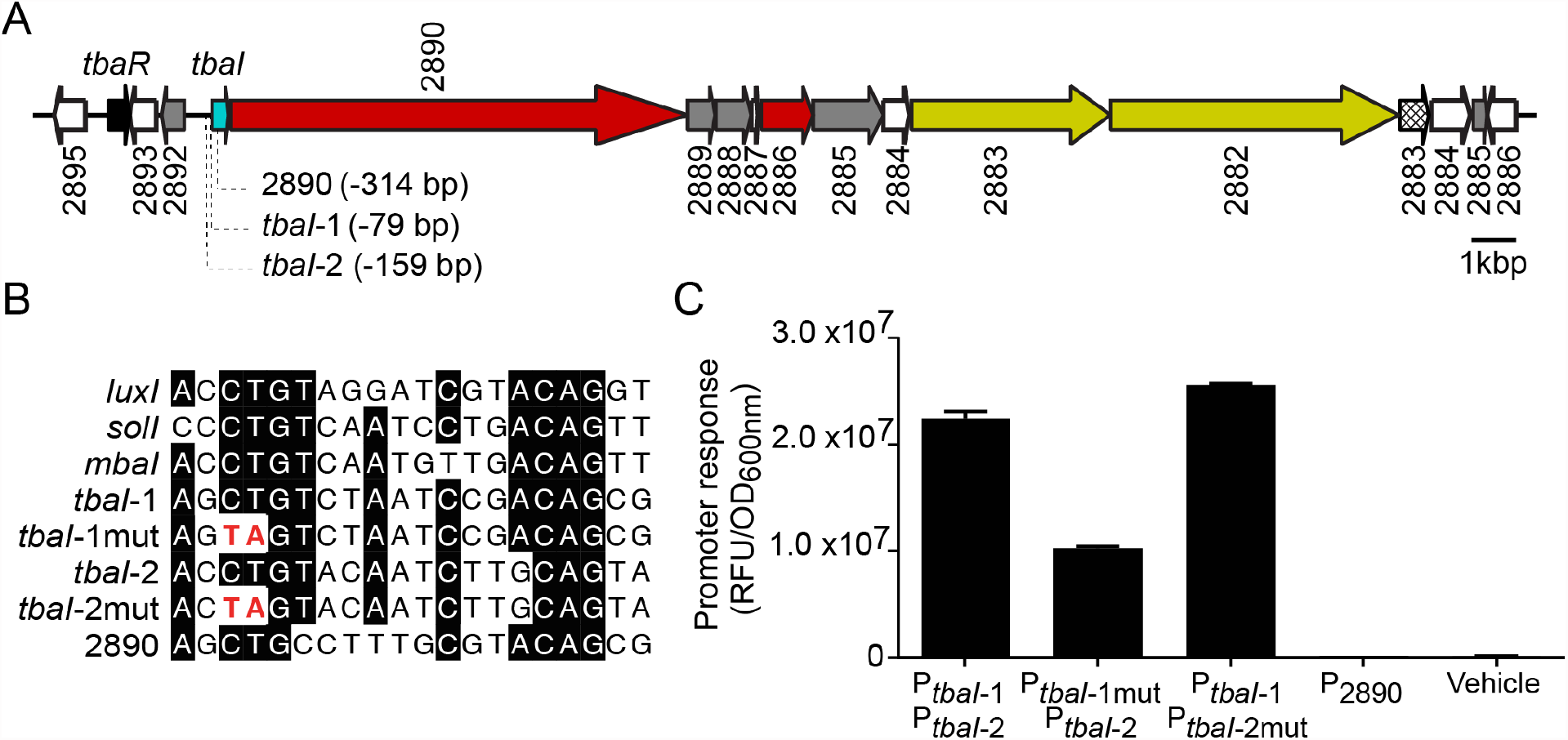
A quorum sensing system in the *Teredinibacter* sp. strain 2052S genome is adjacent to a conserved biosynthetic gene cluster. (A) Quorum sensing genes (*tbaI* and *tbaR*) neighbor a predicted hybrid *trans-*AT-PKS-NRPS biosynthetic gene cluster (3). Identified putative TbaR-binding sites are represented by dotted lines from their position in the cluster, and list the number of base pairs they are located upstream of the start codon of the indicated gene. Numbers correspond to locus tags (K256DRAFT_XXXX) in the Joint Genome Institute Integrated Microbial Genomes (IMG) system (14). Genes are colored as follows according to predicted function in antiSMASH 6.0 (15): *luxI*-family acyl-HSL synthase gene *tbaI*, cyan; *luxR*-family transcription factor gene *tbaR*, black; polyketide synthase genes, red; nonribosomal peptide synthetase genes, yellow; other biosynthetic gene, gray; efflux pump gene, cross-hatched; other genes, white. (B) Comparison of putative TbaR-binding sites upstream of the acyl-HSL synthase gene *tbaI* and PKS gene 2890 with known LuxR-type binding sites in the promoter sequences of *Aliivibrio fischeri luxI, Ralstonia solanacearum solI*, and *Methylobacter tundripaludum mbaI*. (C) Response of *E. coli* reporter strains containing *gfp* fused to different promoter regions with putative TbaR-binding sites shown in 1B to 100 nM C_10_-HSL or ethyl acetate (vehicle). Data are the mean ± standard deviation of three technical replicates and are representative of two independent experiments.

We also found QS genes adjacent to BGCs in the genomes of other shipworm symbionts **(Figure S2)**. However, these BGCs had low similarity to GCF_3, suggesting that QS may regulate the production of other secondary metabolites in these strains. Notably, *Teredinibacter turnerae*, the most well-characterized shipworm symbiont, does not harbor a *luxI*-family synthase gene or complete *luxR*-family transcription factor gene. We have thus far only identified predicted QS systems in isolates from wood-boring shipworms that are not dominated by *T. turnerae* (3).

### 2052S produces and responds to the quorum sensing signal *N*-decanoyl-*L*-homoserine lactone

We characterized the 2052S QS system by first determining if this strain can produce and respond to a QS signal under laboratory conditions. We identified two potential LuxR-family binding sites upstream of the *tbaI* acyl-HSL synthase gene **(Figures 1A, 1B)**, which is often positively autoregulated by its cognate LuxR-family transcription factor upon signal binding (6). We then constructed a two-plasmid reporter system (P_*tbaI*_*-gfp*) in *Escherichia coli* in which one plasmid expresses *tbaR* under its native promoter and the other plasmid contains the *tbaI* promoter, which includes the putative LuxR-family binding sites, fused to *gfp*. Adding organic extract of supernatant from a wild-type (WT) 2052S culture to the P_*tbaI*_*-gfp* reporter strain resulted in a significant increase in GFP fluorescence compared to a solvent control **(Figure S3)**. This confirms that 2052S produces a QS signal and that the LuxR-family homolog, TbaR, binds this signal and activates *tbaI* expression in a positive feedback loop.

We then determined which of the two putative binding sites is primarily used by TbaR by constructing two separate reporter strains containing a CT to TA mutation in the conserved region of each site **(Figure 1B)**. We found that GFP fluorescence was unaffected when the mutation was introduced in P_*tbaI*-2_, suggesting that P_*tbaI*-1_ is the primary TbaR binding site **(Figure 1C)**. However, mutating P_*tbaI-1*_ did not completely abolish *gfp* activation, suggesting that TbaR can also bind P_*tbaI*-2_ in the absence of P_*tbaI*-1_.

In order to isolate and characterize the acyl-HSL signal produced by 2052S, we separated organic supernatant extract by HPLC and tested each fraction using the P_*tbaI*_*-gfp* reporter strain. This resulted in one peak of GFP fluorescence in two adjacent fractions, which was not present in supernatant from an unmarked, in-frame Δ*tbaI* mutant we constructed using sucrose counterselection **(Figure 2A)**. We detected a feature with an *m/z* of 256 in the pooled active fractions using LC-MS, which is consistent with the protonated mass of *N*-decanoyl-*L*-homoserine lactone (C_10_-HSL).

**Figure 2.**
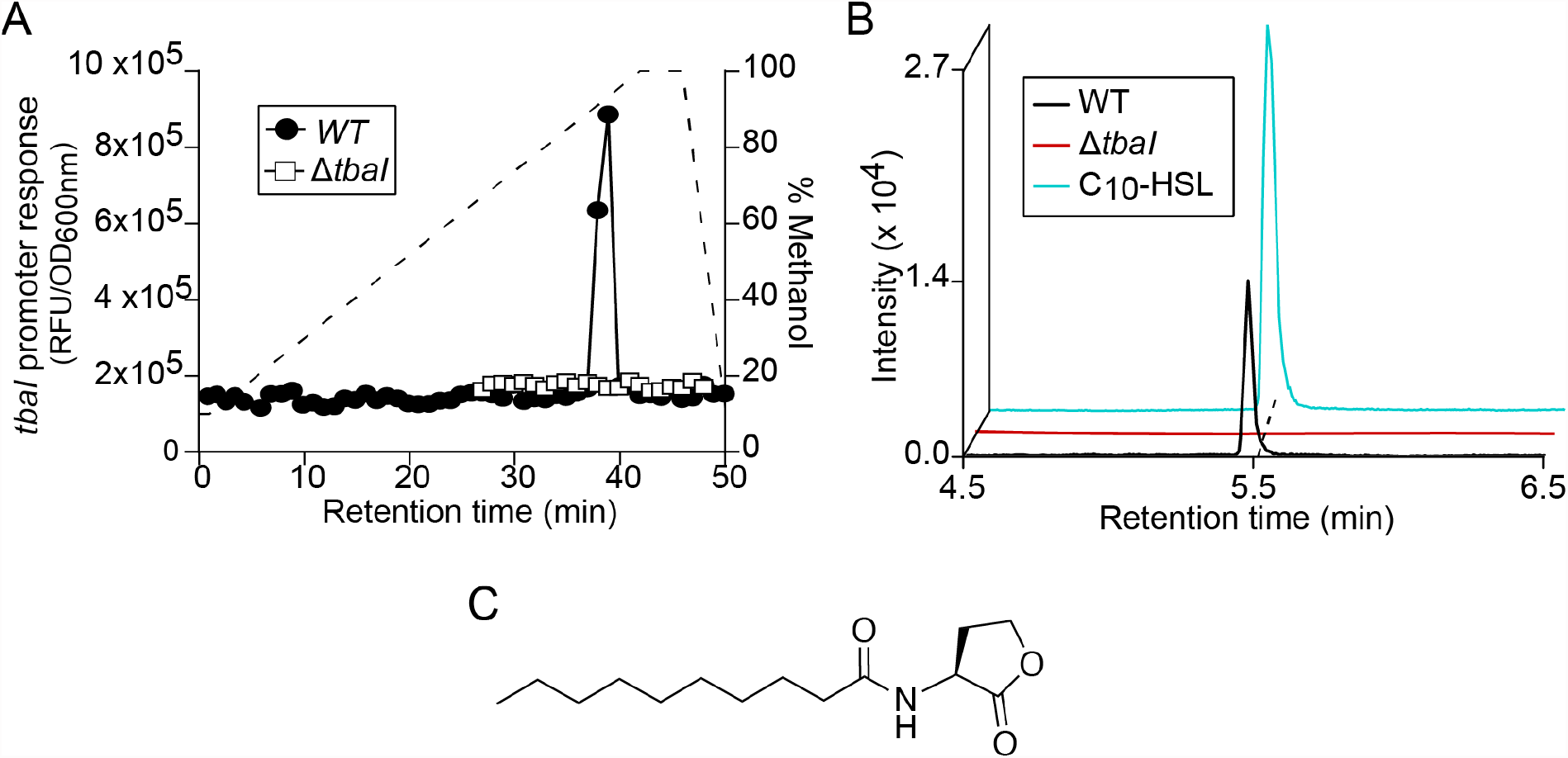
*Teredinibacter* sp. strain 2052S produces the quorum sensing signal C_10_-HSL. (A) P_*tbaI*_-*gfp* activity of HPLC-fractionated culture supernatant extracts from wild-type and *ΔtbaI* strains of 2052S. The dashed line shows the methanol gradient. (B) Extracted ion chromatogram of supernatant extracts of wild-type and *ΔtbaI* strains of 2052S compared to commercial C_10_-HSL signal for *m/z* 279.1812, corresponding to the sodium adduct of C_10_-HSL. Mass tolerance < 5 ppm. (C) Structure of C_10_-HSL.

We confirmed that 2052S produces C_10_-HSL by using high resolution LC-MS/MS to compare organic supernatant extracts from the WT and Δ*tbaI* mutant strains with a commercial standard of C_10_-HSL **(Figure 2B)**. The retention time and fragmentation pattern of the signal produced by WT 2052S were indistinguishable from the commercial standard **(Figure 2B, Table S2)**. Furthermore, the P_*tbaI*_*-gfp* reporter strain was found to be responsive to the commercial standard **(Figure S4)**, and we used the P_*tbaI*_*-gfp* reporter assay to determine that 2052S produces approximately 250 ± 22 nM signal during early stationary phase (OD_600nm_ = 1.2, 32 hours). Together, these results demonstrate that the bacterial endosymbiont 2052S produces and responds to the quorum sensing signal C_10_-HSL.

### Transcription of the conserved biosynthetic gene cluster GCF_3 is regulated by quorum sensing in 2052S

We next sought to determine what 2052S regulates using QS. The Δ*tbaI* mutant is still capable of using cellulose as its primary carbon source, indicating that cellulase production is not regulated by QS in 2052S (all experiments used cellulose as the carbon source, see also **Figure S5)**. We noticed a significant change in the pigmentation of the Δ*tbaI* culture, which we could partially complement by adding exogenous C_10_-HSL **(Figure 3A)**. This suggests that QS may play a role in regulating secondary metabolite production in 2052S. We used RT-qPCR to determine if the conserved GCF_3 BGC adjacent to the QS genes in the 2052S genome is regulated by QS. We quantified the transcription of core genes in this cluster (K256DRAFT_2890, K256DRAFT_2886, and K256DRAFT_2881) in the WT, Δ*tbaI* mutant, and Δ*tbaI* mutant chemically complemented with C_10_-HSL. All three core biosynthetic genes were found to be expressed in a QS signal-dependent manner in a late log-phase culture (OD_600nm_ = 0.8, 24 hours) **(Figure 3B)**.

**Figure 3.**
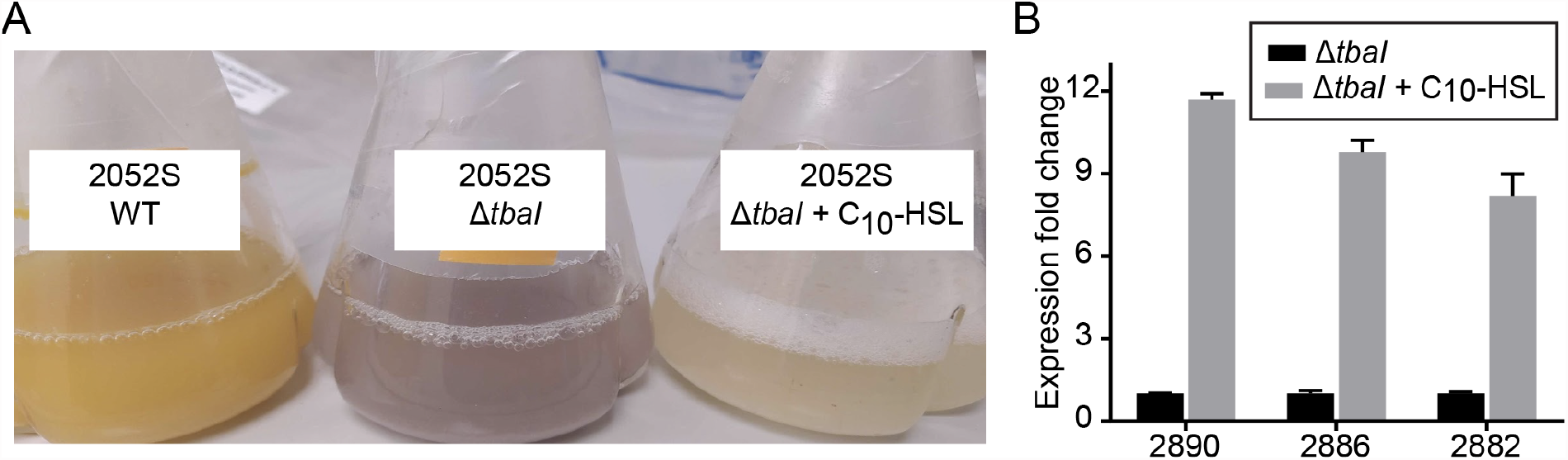
The conserved BGC is transcriptionally activated by the addition of C_10_-HSL in the *ΔtbaI* mutant. (A) Pigmentation phenotype of wild-type (left) and *ΔtbaI* (middle) cultures, and partial reversion to wild-type phenotype upon addition of QS signal to *ΔtbaI* culture (right). (B) RT-qPCR results showing relative expression of GCF_3 genes K256DRAFT_2890, 2886, and 2882 upon addition of C_10_-HSL to the *ΔtbaI* strain normalized to *ΔtbaI* expression in the absence of signal. Data are the mean ± standard deviation of three technical replicates and are representative of two independent experiments.

The first gene in the GCF_3 cluster is K256DRAFT_2890, which encodes a predicted multidomain *trans*-AT-PKS. We identified a putative TbaR-binding site upstream of K256DRAFT_2890 within the *tbaI* gene **(Figures 1A, 1B)**. However, when we created a reporter strain containing this upstream region, it did not drive *gfp* expression in response to C_10_-HSL **(Figure 1C)**. This suggests that both *tbaI* and K256DRAFT_2890 are transcribed together in the same operon, which we confirmed by RT-PCR **(Figure S6)**. Together, these results demonstrate that 2052S uses QS to coordinate the activation of the conserved GCF_3 BGC.

### QS regulates the majority of the extracellular metabolome of 2052S

Because GCF_3 contains predicted efflux pumps **(Figure 1A)**, we sought to identify changes in the extracellular metabolome of 2052S that are controlled by QS in order to determine which secondary metabolites are linked to this cluster. We first constructed an unmarked, in-frame deletion mutant of the 1.3kb polyketide synthase gene K256DRAFT_2886 (Δ2886), and also complemented this mutant using a plasmid containing K256DRAFT_2886 driven by the 2052S *rpoD* promoter (pAWP275). We then used untargeted metabolomics to compare crude organic supernatant extracts from stationary phase cultures of WT 2052S, the *ΔtbaI* mutant, the *ΔtbaI* mutant supplemented with C_10_-HSL, the Δ2886 mutant, and the Δ2886 mutant complemented with pAWP275.

We analyzed the data using MS/MS-based molecular networking within the Global Natural Products Social molecular networking (GNPS) platform, which clusters metabolites based on MS/MS fragmentation patterns (13). This analysis produced a primary dataset of 442 features from raw LC-MS/MS scans. We further refined this dataset by removing features found in the uninoculated growth medium as well as those not found in both independent replicates, resulting in a final dataset of 256 features **(Figure 4A)**. The majority (175/256, 68%) of features were only detectable in the supernatant of QS-active strains **(Figure 4B)**. This indicates that QS plays an important role in the regulation of secondary metabolism in this shipworm endosymbiont. Notably, none of the 2052S extracellular secondary metabolites had matches to compounds in the GNPS spectrum library. This highlights the potentially unique biosynthetic potential of shipworm endosymbionts.

**Figure 4.**
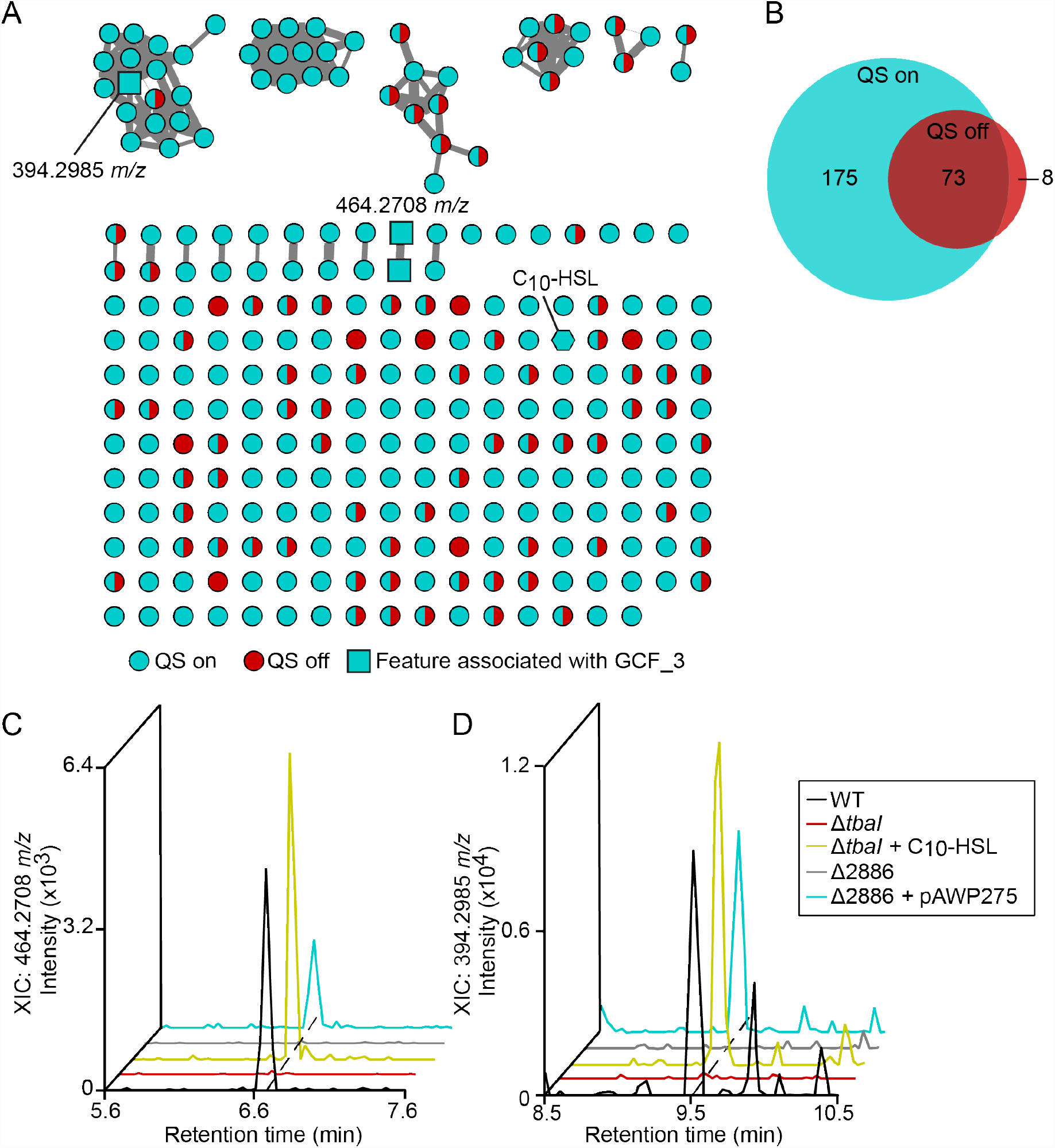
Quorum sensing regulates the majority of extracellular metabolites produced by 2052S. (A) Molecular network of untargeted metabolomics data of supernatant extracts from cultures of WT, *ΔtbaI*, and *ΔtbaI* supplemented with C_10_-HSL. Features found in samples where QS is on (WT, Δ*tbaI* + C_10_-HSL) are shown as cyan nodes, features found in the Δ*tbaI* culture, where QS is off, are shown as red nodes, and features found in both samples where QS is on and off are shown as split cyan and red nodes. Features identified as associated with GCF_3 (present in WT, Δ*tbaI* + C_10_-HSL, and Δ2886 +pAWP275, but absent in Δ*tbaI* and Δ2886) are shown as square nodes, and C_10_-HSL is shown as a hexagonal node. Edge width is scaled with cosine similarity score. (B) Venn diagram of features associated with the QS regulon. (C,) Extracted ion chromatogram of *m/z* 464.2708 in culture supernatant extracts. Mass tolerance < 5 ppm. (D) Extracted ion chromatogram of *m/z* 394.2985 in culture supernatant extracts. Mass tolerance < 5 ppm.

To identify the putative product of the GCF_3 BGC in 2052S, we focused on extracellular metabolites present in cultures of the WT, the *ΔtbaI* mutant supplemented with C_10_-HSL, and the Δ2886 mutant complemented with pAWP275, but absent in the Δ*tbaI* and Δ2886 mutants. We identified two putative metabolites matching this pattern that may be products of this gene cluster **(Figures 4C, 4D)**, including a feature with a precursor ion mass of 394.2985 *m/z* found in the largest cluster in the network **(Figure 4A)**. This cluster was found almost exclusively in QS-active samples. These metabolites will require further investigation to determine their structure and function.

We have identified and characterized a QS system in *Teredinibacter* sp. strain 2052S, a symbiont of the wood-boring shipworm *B*. cf. *thoracites*. We determined that 2052S produces and responds to the signal C_10_-HSL, and that this signal regulates the activation of a BGC that is conserved among all wood-boring shipworm symbiont isolates, termed GCF_3. It is possible that secondary metabolites produced by shipworm endosymbionts play a role in establishing and maintaining the relationship between these bacteria and their host. The discovery of a symbiont that regulates its extracellular secondary metabolism using QS is consistent with this hypothesis, as QS is often thought to enable bacterial symbionts to differentiate between planktonic and host-associated states. More studies will be needed to understand the role of these metabolites, as well as QS, in this symbiotic relationship.

## MATERIALS AND METHODS

### Plasmid Construction

All plasmids were constructed using Gibson assembly (16), with the exception of the reporter plasmids pAWP239, pAWP381, pAWP479, and pAWP480, which were constructed by inserting upstream gene sequences into the promoter probe plasmid pPROBE-GFP[LVA] (17) at the EcoRI and SacI restriction sites. The upstream sequences in pAWP479 and pAWP480, which contain CT-to-TA mutations, were ordered as gBlocks from Integrated DNA Technologies. Plasmids and primers used in this study are listed in Table 1 and Table 2, respectively.

**Table 1.**
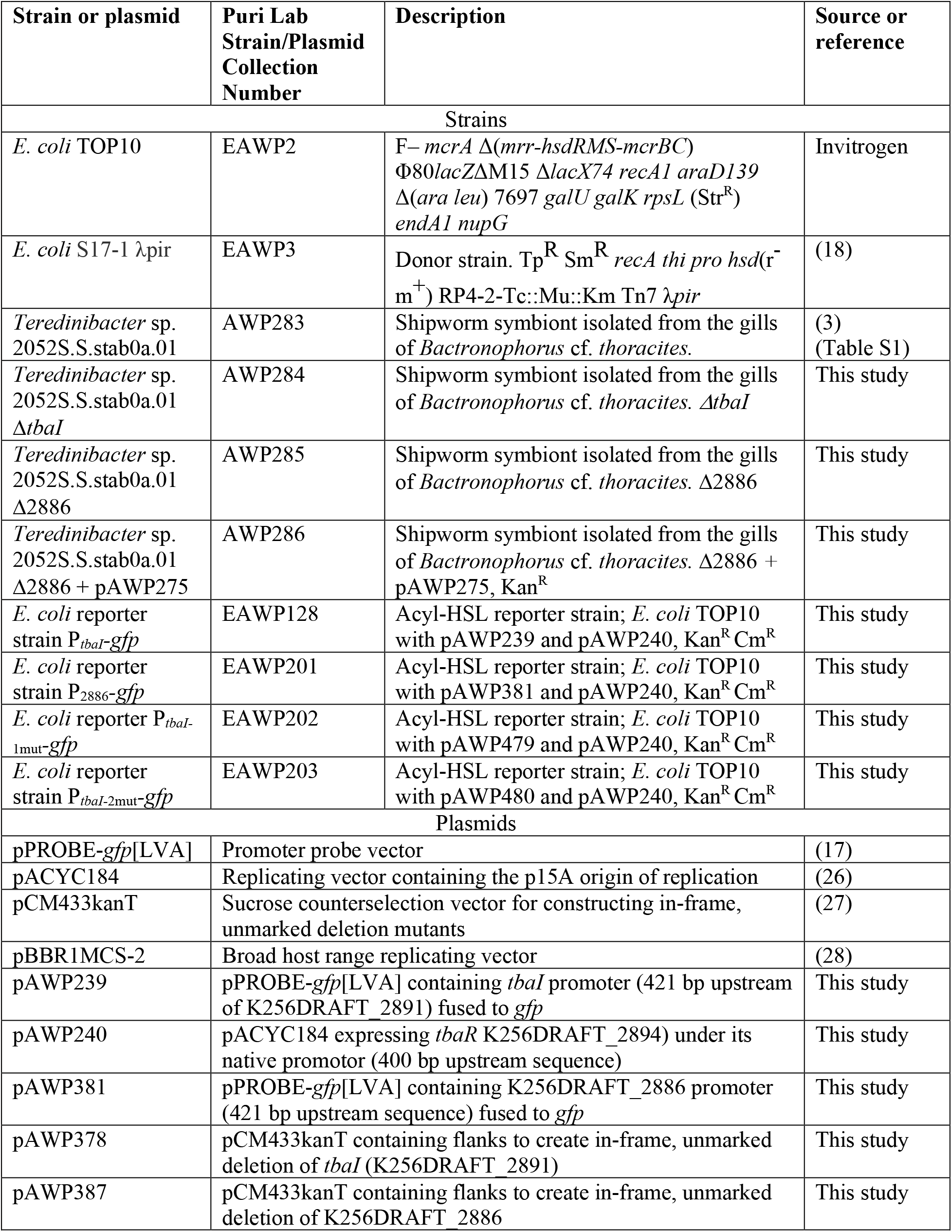
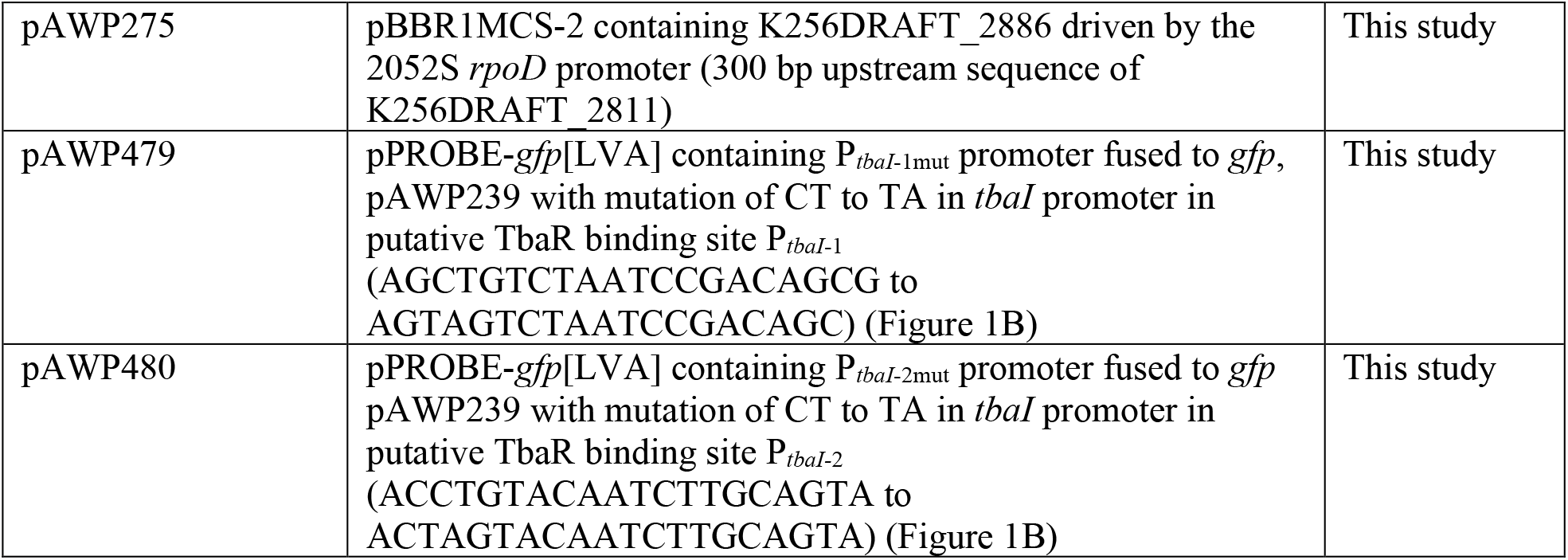
Strains and plasmids used in this study.

**Table 2.**
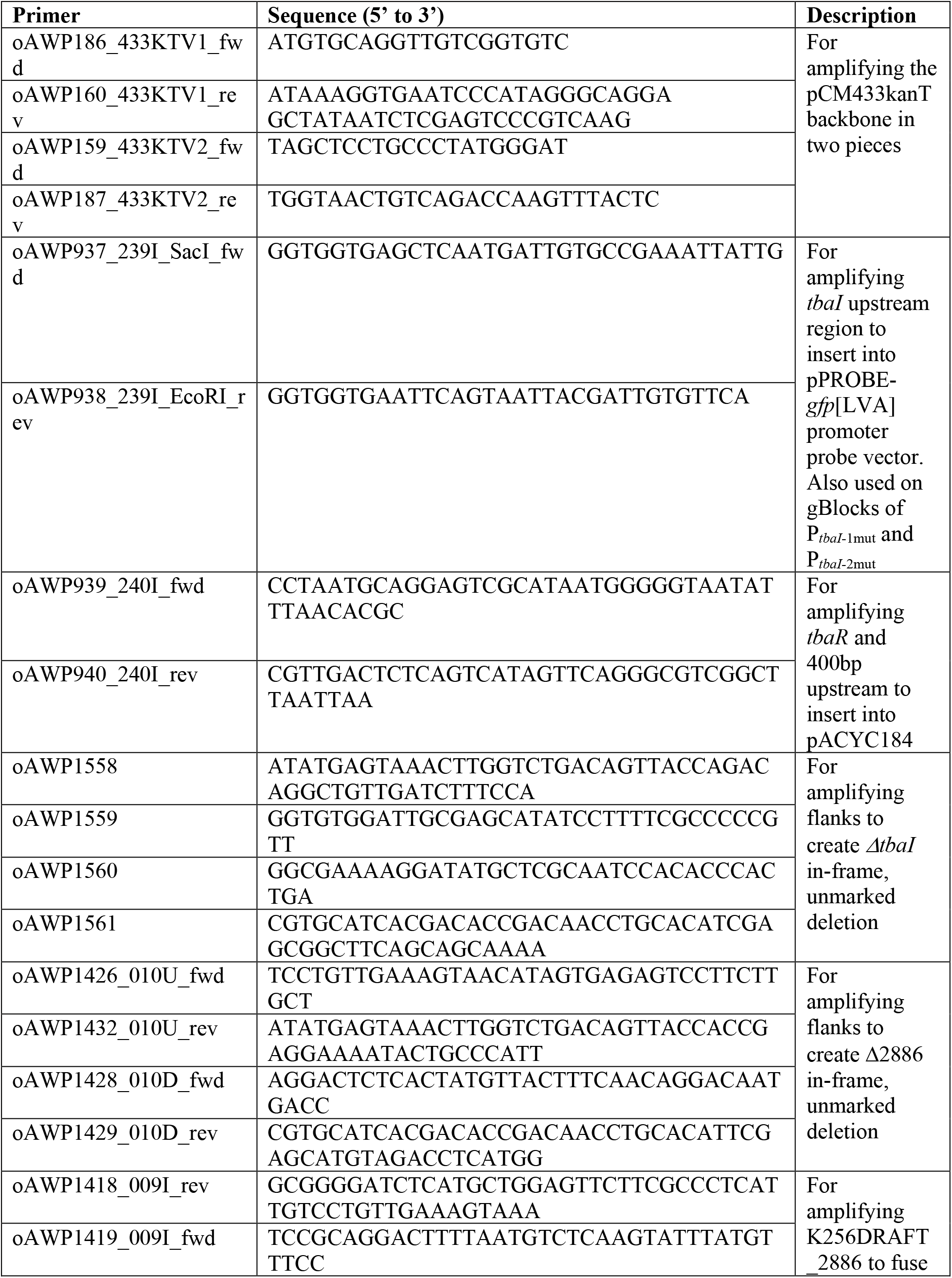
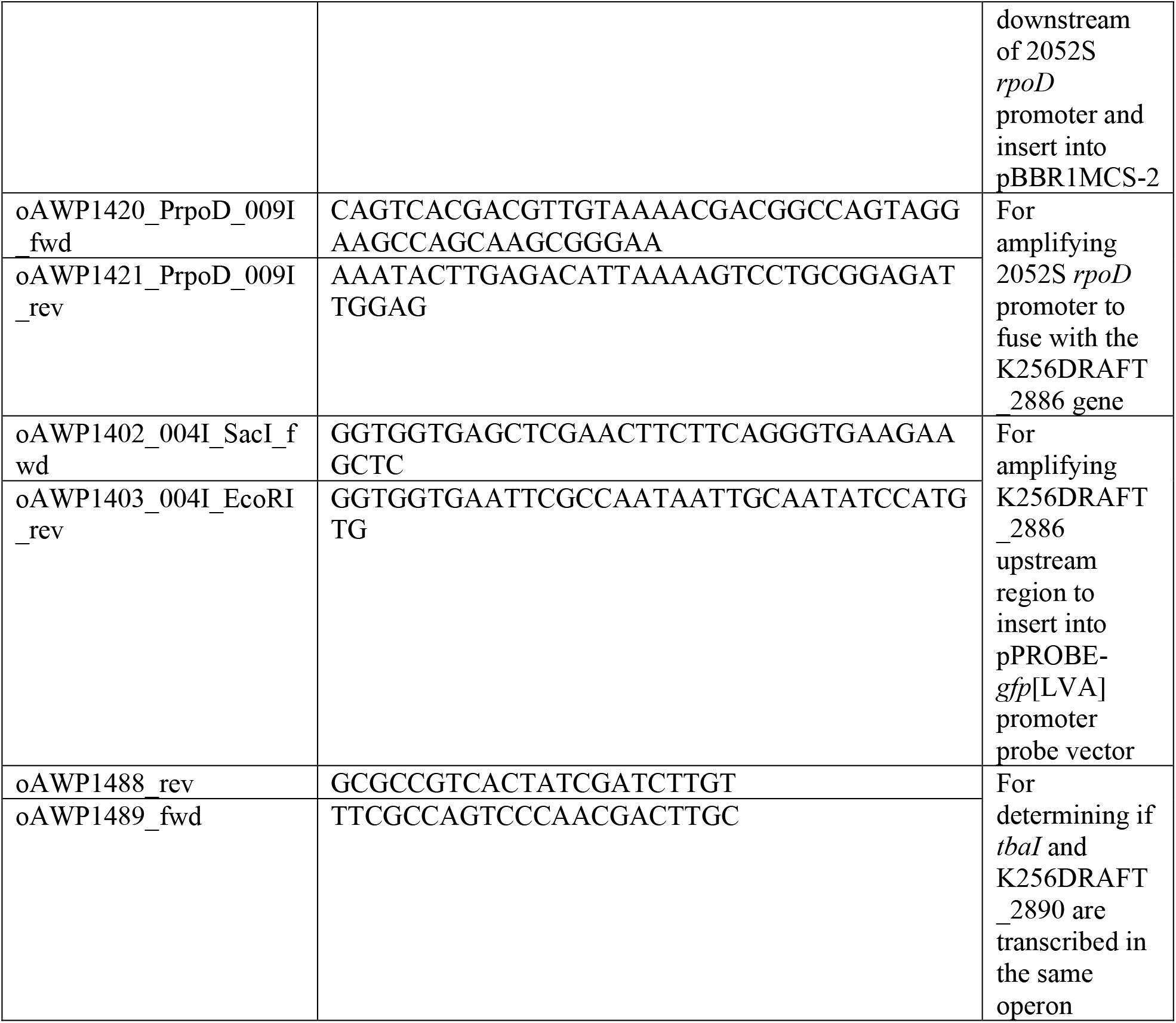
Cloning and diagnostic primers used in this study.

### Strain Growth

Strains used in this study are listed in Table 1. *E. coli* strains were grown in lysogeny broth (LB) at 37°C. *Teredinibacter* sp. strain PMS-2052S.S.stab0a.01 (2052S) was isolated from *Bactronophorus cf. thoracites* (PMS-1959H) collected from Butuan, Agusan del Norte, Philippines (Table S1) and was grown on shipworm basal media (30°C, 200 rpm) as previously described by Waterbury *et al*. (1983) (2). SBM contains NaCl (19.8 g/L), NH_4_Cl (267.5 mg/L), MgCl_2_·6H_2_O (8.95 g/L), Na_2_SO_4_ (3.31 g/L), CaCl_2_·2H_2_O (1.25 g/L), NaHCO_3_ (0.162 g/L), Na_2_CO_3_ (10 mg/L), KCl (0.552 mg/L), KBr (81 mg/L), H_3_BO_3_ (21.5 mg/L), SrCl_2_·6H_2_O (19.8 mg/L), KH_2_PO_4_ (3.82 mg/L), NaF (2.48 mg/L), Na_2_MoO_4_·2H_2_O (2.5 mg/L), MnCl_2_·4H_2_O (1.8 mg/L), ZnSO_4_. 7H_2_O (0.22 mg/L), CuSO_4_·5H_2_O (0.079 mg/L), Co(NO_3_)_2_·6H_2_O (0.049 mg/L), Fe-EDTA complex (4.15 mg/L), and HEPES (4.76 g/L) adjusted to pH = 8.0. The carbon source used was Sigmacell cellulose type 101 (0.2 g/L).

### Genetic Manipulation

Genetic manipulation of all strains derived from wild-type 2052S were performed at 30ºC. Verified plasmids were conjugated into 2052S using the *E. coli* donor strain S17-1 (18) using the following method. 500 µL of exponentially growing cultures of donor and recipient strains were pelleted and washed with sterile ultrapure water. The two pellets were then combined in a total volume of 50 µL and spotted onto an SBM plate containing 10% (v/v) nutrient broth, and subsequently incubated for two days. Successful conjugants were selected on SBM plates containing kanamycin (50 µg mL^-1^). To construct the unmarked deletion mutants Δ*tbaI* and Δ2886, kanamycin-resistant integrants (single crossovers) were restreaked and grown in SBM broth with no kanamycin before being spread plated onto an SBM plate containing 5% (v/v) sucrose for counterselection. The resulting colonies were screened for double crossovers by kanamycin sensitivity and colony PCR before the final mutant was verified by PCR and Sanger sequencing.

### Identification of LuxR-type binding site

Nucleotide sequences were identified as putative LuxR-type binding sites if they satisfied the following criteria: (1) located within 400 bp upstream of the translational start site, (2) matched the general NNCTG-N_10_-CAGNN pattern with one mismatch or less, and (3) contained eight or more base pairs with dyad symmetry (10, 19).

### Acyl-HSL reporter assay

The reporter assay was performed as described in (20). Briefly, cultures were centrifuged at 16,000 g and the supernatant from a wild-type (WT) culture of 2052S was extracted twice with an equal volume of ethyl acetate containing 0.01% (v/v) acetic acid. The organic phase was subsequently dried via nitrogen stream. The dried extract was then resuspended in acidified ethyl acetate, aliquoted into 1.5 mL tubes, and before being re-dried before an overnight *E. coli* reporter strain diluted to an OD of 0.1 was added to the tube. After four hours of incubation (37C, 200 rpm), GFP fluorescence (485-nm excitation, 510-nm emission) and absorbance at 600 nm were measured in a 96-well black, clear-bottomed plate by using a plate reader (SpectraMax i3x).

### Isolation and characterization of the QS signal

The acyl-HSL signal produced by 2052S was extracted from supernatant of a 50 mL culture extracted at early stationary phase (OD_600nm_ = 1.2, 32 hours) as described above. The supernatant extract was resuspended in methanol and 20% (from 10 mL of culture) was separated by HPLC using a Waters SunFire C_18_ column (4.6 × 100 mm, 5µm) at 1.0 ml/min using a linear gradient of 10% to 100% methanol in water over 50 minutes. A 5-μl aliquot of each 1.0-ml fraction was analyzed using the *E. coli* reporter strain described above. The pooled adjacent fractions that showed GFP activity were then analyzed using high-resolution tandem mass spectrometry. Confirmation of the identified signal was performed using a C_10_-HSL standard purchased from Cayman Chemical.

### RNA preparation

Exponentially growing cultures of 2052S were diluted to an OD_600nm_ of 0.01 and were grown until log-phase (OD_600nm_ of 0.8, 24 hours) prior to RNA extraction. For *ΔtbaI* + C_10_-HSL signal, 2 mM C_10_-HSL in DMSO was added to a final concentration of 2 μM every 12 hours. Subsequently, cultures were chilled on ice and then centrifuged at 4700 rpm for 15 min at 4°C. Pellets were then stored at −80°C until further processing. Cell pellets were lysed by bead beating with 0.1-mm zirconia-silica beads in 1 ml TRIzol (ThermoFisher). 200 µL of chloroform was then added and the mixture was separated by centrifugation using phasemaker tubes (ThermoFisher). Subsequently 1.5 volumes of 100% ethanol was added to the aqueous phase of the extract, which was then used for DNase I treatment (Invitrogen) and cleanup using an Invitrogen RNA PureLink mini kit according to the manufacturer’s instructions. The resulting purified RNA was checked for DNA contamination by Nanodrop and PCR using the degenerate 16S primers 27F and 1492R.

### RT-qPCR

cDNA was prepared for RT-qPCR by reverse transcribing one microgram of the extracted RNA using iScript Reverse Transcription Supermix (BioRad). qPCR was performed using iTaq-Universal SYBR Green Supermix (BioRad) containing 400 nM primers and cDNA normalized across all samples in a total volume of 10 µL. qPCR reactions were performed on a BioRad CFX Opus 96 thermal cycler, and threshold cycle (C_T_) values were calculated using BioRad CFX Maestro software. All primers and their corresponding gene targets are listed in Table 3.

**Table 3.**
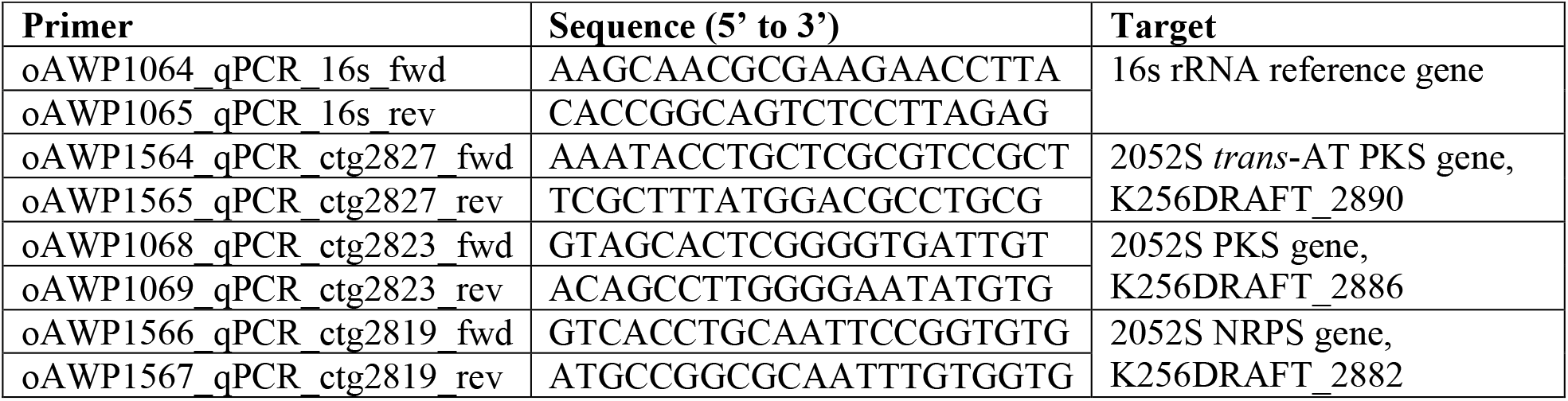
Reverse transcription quantitative PCR primers used in this study.

### High-resolution LC-MS/MS for acyl-HSL identification

Mass spectrometry data were collected using a Waters Acquity I-class ultra-high pressure liquid chromatograph coupled to a Waters Xevo G2-S quadrupole time-of-flight mass spectrometer. An Acquity UPLC BEH C_18_ column (2.1 × 50 mm) was used for separation of samples. Solvent A: Water + 0.1 % (v/v) formic acid, Solvent B: Acetonitrile + 0.1% (v/v) formic acid. The sample was eluted from the column using a ten-minute linear solvent gradient: 0-0.1 min, 1% B; 0.1 - 10 min, 100% B. The solvent flow rate was 0.45 mL min-1. Mass spectra were collected in positive ion mode, with following parameters: 3 kV capillary voltage; 25 V sampling cone voltage; 150 °C source temperature; 500 °C desolvation temperature; nitrogen desolvation at 800 L/hr. The fragmentation spectra were collected using the same parameters with a 10-25 eV collision energy ramp. The lockspray solution was 200 pg/μL leucine enkephalin. The lockspray flow rate was 6 μL/min. Sodium formate was used to calibrate the mass spectrometer.

### Untargeted high-resolution LC-MS/MS for molecular networking

HR-MS/MS data was obtained from 100% methanol elution of HP-20 Diaion resin (Sigma) incubated with culture supernatant for at least 4 hours collected from a late log phase (48 hours) culture of WT, Δ*tbaI*, Δ2886, Δ2886 *+* pAWP275 and Δ*tbaI +* C_10_-HSL (2 μM supplemented every 12 hours). Samples were passed through an C_18_ solid phase extraction cartridge prior to dissolving in 50% ACN/H2O at a final concentration of 1.0 mg/mL. For data collection, the suggested settings for Waters mass spectrometer in (21) were used. An Acquity UPLC BEH C_18_ column (2.1 × 50 mm) was used for separation of samples. Solvent A: Water + 0.1 % (v/v) formic acid, Solvent B: Acetonitrile + 0.1% (v/v) formic acid. A flow rate of 0.6 mL/min was used with the following gradient: 1%–100% B, 0–12 min; 100% B, 12–13 min; and a column reconditioning phase until 15 min. The following parameters were used: 2.5 kV capillary voltage; 20 V sampling cone voltage; 120 °C source temperature; 350 °C desolvation temperature; desolvation gas flow at 800 L/hr. MS^1^ acquisition range was set to *m/z* 100 - 1,500 with a scan time of 0.1 s in data-dependent acquisition mode. The top 5 most abundant MS^1^ ions were selected in each scan and up to five MS^2^ scans in CID mode was acquired with 0.1 s scan time in positive mode. MS survey was set to switch to MS2 acquisition when TIC rises above an intensity of 5.0 × 10^3^, and MS^2^ acquisition switch back to MS survey after 0.25 s have elapsed. The collision energy gradient was set to gradient parameters for as follows: 20 to 40 V for 100 Da to 60 to 80 V for 1500 Da.

### Molecular Networking

A molecular network was created using the online workflow (https://ccms-ucsd.github.io/GNPSDocumentation/) on the GNPS website (http://gnps.ucsd.edu). The data was filtered by removing all MS/MS fragment ions within ± 17 Da of the precursor m/z. MS/MS spectra were window filtered by choosing only the top 6 fragment ions in the ± 50 Da window throughout the spectrum. The precursor ion mass tolerance was set to 0.02 Da and a MS/MS fragment ion tolerance of 0.02 Da. A network was then created where edges were filtered to have a cosine score above 0.7 and more than 6 matched peaks. Further, edges between two nodes were kept in the network if and only if each of the nodes appeared in each other’s respective top 10 most similar nodes. Finally, the maximum size of a molecular family was set to 100, and the lowest scoring edges were removed from molecular families until the molecular family size was below this threshold. The spectra in the network were then searched against GNPS spectral libraries (22, 23). The library spectra were filtered in the same manner as the input data. All matches kept between network spectra and library spectra were required to have a score above 0.7 and at least 6 matched peaks. The DEREPLICATOR was used to annotate MS/MS spectra (24). The molecular networks were visualized using Cytoscape software (25).

## Data Deposition and Job Accessibility

The mass spectrometry data were deposited in the public repository MassIVE: https://massive.ucsd.edu/ProteoSAFe/dataset.jsp?task=f823719f2c5f4fca903bfe49f6964f45.

The molecular networking job can be publicly accessed at: https://gnps.ucsd.edu/ProteoSAFe/status.jsp?task=d4ce2cc7d4db423eb00a4d2876993b50.

## AUTHOR CONTIBUTIONS

JMDR and AWP designed experiments. JMDR and EGM performed experiments. MAA isolated 2052S. GPC, MGH, and AWP oversaw and supported the research. JMDR and AWP wrote the manuscript. All authors have read and approved of the final version of the manuscript.

## COMPLIANCE

The work was completed under supervision of the Department of Agriculture-Bureau of Fisheries and Aquatic Resources, Philippines (DA-BFAR), in compliance with all required legal instruments and regulatory issuances covering the conduct of the research.

## ACKNOWLEDGEMENTS

We thank D. Petras (University of Tübingen) for reading the manuscript and providing helpful advice on the molecular networking analysis. We thank H. Naka (University of Utah) for initial help with *T. turnerae* genetics. This work was supported by National Institutes of Health grant R00 GM118762 (to AWP) and National Institutes of Health Fogarty International Center Philippine Mollusk Symbiont-International Cooperative Biodiversity Group (PMS-ICBG) grant U19TW008163 (to GPC and MGH). This work was also supported by funding from the Undergraduate Research Opportunities Program at the University of Utah (to EGM).

