## Supplementary Figures and Tables for "A conserved biosynthetic gene cluster is regulated by quorum sensing in a shipworm symbiont"

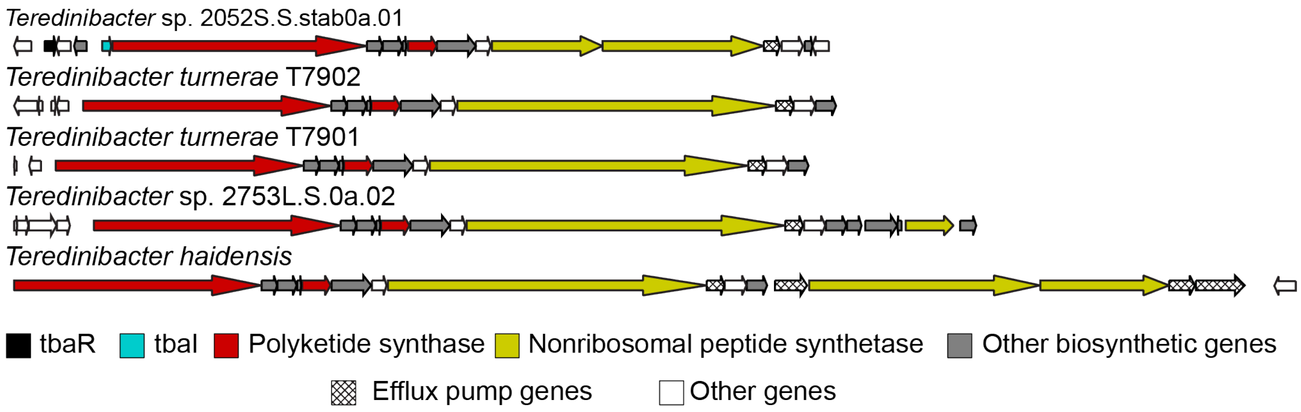


**Figure S1.** GCF_3 is conserved in cellulolytic shipworm symbionts. Representative shipworm symbionts and their biosynthetic gene clusters belonging to GCF_3. Amino acid identity of core biosynthetic genes is at least 65% in all cases. Genes are colored according to predicted function in antiSMASH 6.0 (1).

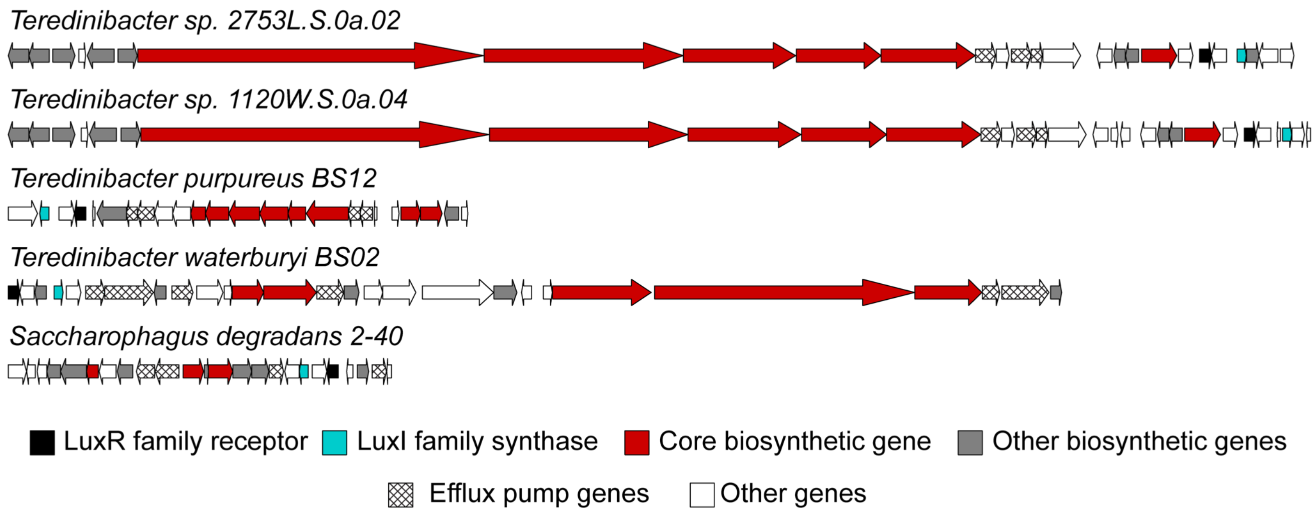


**Figure S2.** Biosynthetic gene clusters co-located with quorum sensing genes in other shipworm symbiont genomes. Genes are colored according to predicted function in antiSMASH 6.0 (1).

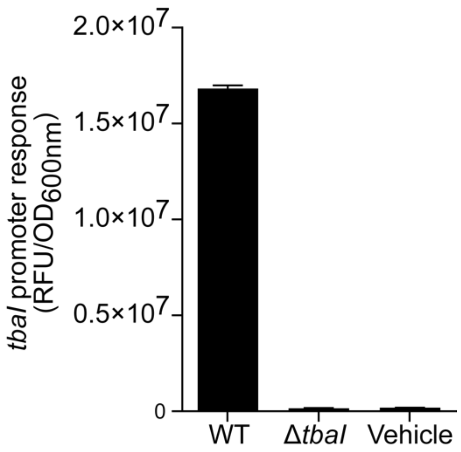


**Figure S3**. Response of $P_{tbaI}$-*gfp E. coli* reporter strain EAWP128 to crude extracts of 2052S wild-type and Δ*tbaI* strains. Vehicle: ethyl acetate.

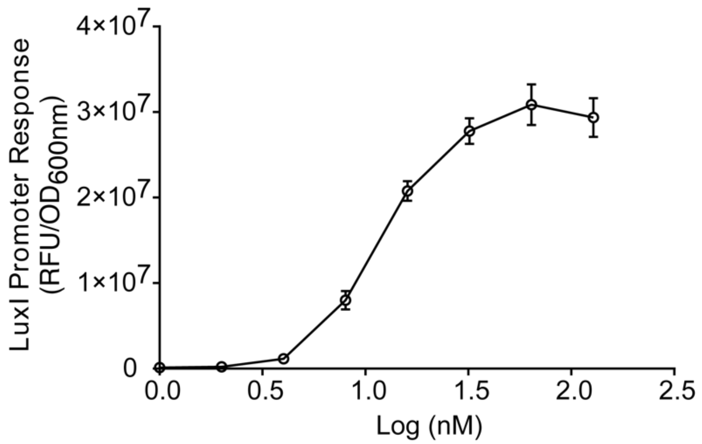


**Figure S4**. Response of $P_{tbaI}$-*gfp E. coli* reporter strain EAWP128 to $C_{10}$-HSL.

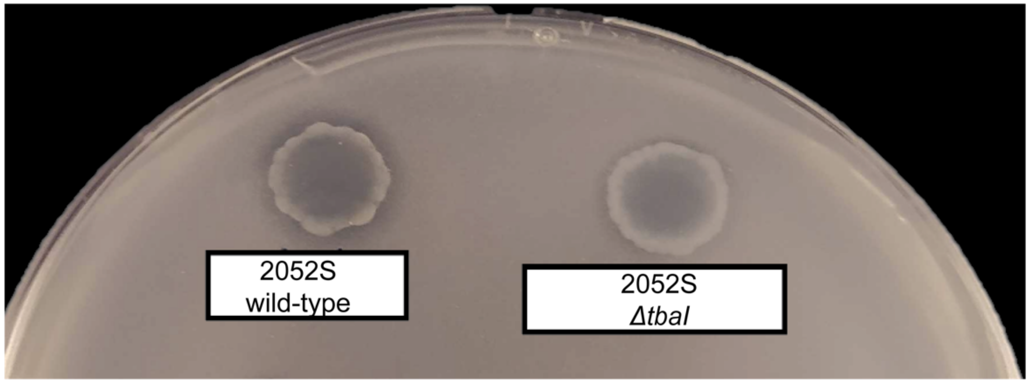


**Figure S5.** The Δ*tbaI* strain retains its cellulolytic ability. Small halo around the spotted colony on SBM cellulose plate indicates cellulolysis.

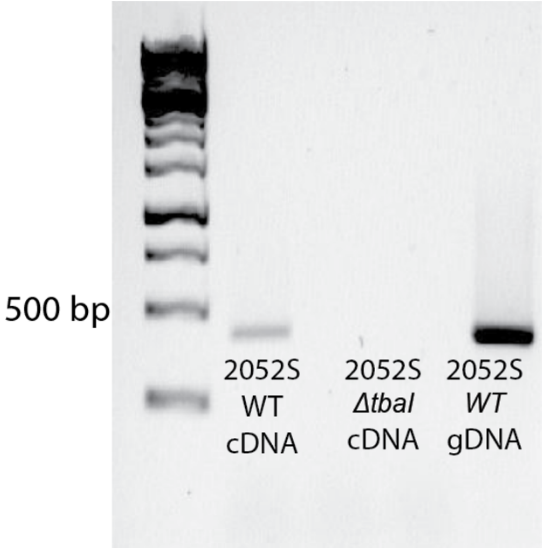


**Figure S6.** The PKS gene K256DRAFT_2890 is co-transcribed with *tbaI*. Primers were designed to span both open reading frames of *tbaI* and K256DRAFT_2890. The PCR template was either genomic DNA (gDNA) or cDNA from wild-type 2052S and Δ*tbaI*. Δ*tbaI* has no band because the primer annealing site in *tbaI* was deleted. Expected product size: 427 bp.

**Table S1.** *Teredinibacter* sp. PMS-2052S.S.stab0a.01 (2052S) strain information

| Isolate name | Metabolic type | Host shipworm | Collection site | Sequencing center | Estimated genome size (bp) | % GC | IMG Genome ID | Reference |
|---|---|---|---|---|---|---|---|---|
| PMS-2052S.S.stab0a.01 | Cellulolytic | PMS-1959H *Bactronophorus* cf. *thoracites* | Butuan, Agusan del Norte, Philippines | JGI-DOE | 5.635,926 | 54.68 | 2541046951 | (2) |

**Table S2.** HR-MS/MS peak list for $C_{10}$-HSL ($[M+H]^+$) produced by 2052s compared to commercial standard. The 10 most intense signals are shown for each sample.

| **2052S** | | **Standard** | |
|---|---|---|---|
| *m/z* | Intensity | *m/z* | Intensity |
| 102.0563 | 6.50E+03 | 102.0565 | 2.90E+04 |
| 256.1914 | 6.30E+03 | 256.1915 | 2.90E+04 |
| 155.143 | 6.10E+03 | 155.1431 | 2.10E+04 |
| 238.181 | 1.20E+03 | 238.1813 | 9.20E+03 |
| 95.0874 | 9.40E+02 | 95.0871 | 8.80E+03 |
| 81.0723 | 8.70E+02 | 81.0718 | 8.60E+03 |
| 74.0631 | 8.60E+02 | 74.0629 | 8.30E+03 |
| 228.194 | 8.10E+02 | 156.1468 | 8.10E+03 |
| 156.1456 | 7.60E+02 | 137.1328 | 8.00E+03 |
| 137.1326 | 7.32E+02 | 228.1968 | 7.90E+03 |
